# Regulation of bacterial stringent response by an evolutionary conserved ribosomal protein L11 methylation

**DOI:** 10.1101/2023.06.29.546327

**Authors:** Hanna E. Walukiewicz, Yuliya Farris, Meagan C. Burnet, Sarah C. Feid, Youngki You, David Christensen, Samuel H. Payne, Alan J. Wolfe, Christopher V. Rao, Ernesto S. Nakayasu

**Author notes:** Corresponding author: Ernesto S. Nakayasu.

## Abstract

Lysine and arginine methylation is an important regulator of enzyme activity and transcription in eukaryotes. However, little is known about this covalent modification in bacteria. In this work, we investigated the role of methylation in bacteria. By reanalyzing a large phyloproteomics dataset from 48 bacterial strains representing 6 phyla, we found that almost a quarter of the bacterial proteome is methylated. Many of these methylated proteins are conserved across diverse bacterial lineages, including those involved in central carbon metabolism and translation. Among the proteins with the most conserved methylation sites is ribosomal protein L11 (bL11). bL11 methylation has been a mystery for five decades, as the deletion of its methyltransferase PrmA causes no cell growth defects. A comparative proteomics analysis combined with a guanosine polyphosphate assay of the Δ*prmA* mutant in *Escherichia coli* revealed that bL11 methylation is important for stringent response signaling. Moreover, we show that the Δ*prmA* mutant has an abnormal polysome profile, suggesting a role in ribosomal homeostasis during stationary growth phase. Overall, our investigation demonstrates that the evolutionary conserved bL11 methylation is important for stringent response signaling and ribosomal homeostasis.

**Importance:** Protein methylation in bacteria was first identified over sixty years ago. Since then, its functional role has been identified for only a few proteins. To better understand the functional role of methylation in bacteria, we analyzed a large phyloproteomics dataset encompassing 48 diverse bacteria. Our analysis revealed that ribosomal proteins are often methylated at conserved residues, suggesting that methylation of these sites may have a functional role in translation. Further analysis revealed that methylation of ribosomal protein L11 is important for stringent response signaling and ribosomal homeostasis.

## Introduction

Proteins can be modified by proteolytic cleavage or the addition/removal of chemical groups. These post-translational modifications (PTMs) regulate diverse cellular processes, such as enzymatic activities, protein-protein interactions, protein localization, and protein stability (1, 2). Like other PTMs, methylation of lysine and arginine residues has been shown to occur in proteins that possess a variety of functions (3, 4). Global proteomic analyses of the methylated proteome have enabled the identification and quantification of many modified sites in cells (5-7). In eukaryotes, the most studied methylated proteins are histones and transcription factors, which regulate chromatin structure and gene expression, respectively (4, 8). However, little is known about the distribution and function of protein methylation in bacteria (2).

Protein methylation in bacteria was first described over sixty years ago for flagellin in *Salmonella enterica* (9). However, the function of flagellin methylation was only identified in 2020, when it was shown to promote bacterial adhesion and mediate host cell invasion (10). Aside from flagellin, the bacterium *Chlamydophila pneumoniae* has a Su(var)3-9, Enhancer-of-zeste and Trithorax (SET) domain-containing methyltransferase that modifies two histone H1-like proteins (11), suggesting that gene expression regulation by methylation is conserved throughout evolution. Lysine tri-methylation sites on the ribosomal protein L11 (bL11, also known as RPL11 or RplK) by the methyltransferase PrmA were found in the bacterial species *Escherichia coli* and *Thermus thermophilus*, and in plant chloroplasts and mitochondria (12, 13). Despite being first described in the early seventies (14), the role of bL11 methylation has remained unknown since its loss does not affect cell growth or protein synthesis (10, 15).

Here, we sought to determine additional roles for protein methylation in bacteria. Based on the hypothesis that evolutionary conserved features are more likely to have important roles (16), we investigated conserved methylation sites in bacteria using a phyloproteomics dataset containing 48 bacterial strains from 6 phyla. Our analysis revealed enrichment of protein methylation in several pathways conserved across evolution, including central carbon metabolism and translation. Among the most conserved sites are the lysine trimethylations of bL11. In bacteria, the bL11 protein performs several functions. It is the target of the antibiotic thiostrepton (17), regulates translation termination (18), and functions as a regulator of the stringent response, which occurs as the culture enters stationary phase (19). In the stringent response, bL11 helps the ribosome sense nutrient scarcity by detecting deacylated (without amino acids) tRNAs, inducing the production of the alarmone guanosine polyphosphate ((p)ppGpp), which in turn signals the cells to halt growth and conserve resources (20). Therefore, we studied the role of L11 methylation by combining proteomics with molecular biology and biochemistry. Our results show that this modification has important roles in stringent response signaling and in maintaining ribosomal homeostasis.

## Material and Methods

### Phyloproteomics data reanalysis

Data from a previously reported bacterial phyloproteomics analysis (21) were reanalyzed for identification of methylation sites. Raw mass spectrometry files were converted to the PSI open format mzML, using msConvert (22). Files were re-calibrated with the mzRefinery tool (23) and searched against species-specific sequence databases (21), using MS-GF+ (24).

Searching parameters were precursor mass tolerance as 20.0 ppm; oxidation of methionine as dynamic modification; maximum modifications per peptide as 3; isotope error as -1,1; and tryptic digestion in both peptide termini with a maximum of 2 mis-cleavage sites. Mono- and di-methylation of both lysine and arginine residues, and tri-methylation of lysine were performed in separated searches. Spectral-peptide matches were filtered with a MS-GF probability score ≤ 1e-10, resulting in a false discovery rate < 1% at both peptide and protein levels.

### Structural analysis

The structure of the 70S *E. coli* ribosome (PDB index 5UYK) was downloaded from Protein Data Bank - PDB and visualized with Discovery Studio Visualizer 4.5 program.

### Multiple sequence alignment

RplK (bL11 gene name) sequences were aligned using UGENE (v 46.0) software with Clustal Omega, with the phylogenetic tree built with the Jones-Taylor-Thornton distance matrix model (25).

### Cell culture

*E. coli* BW25113 and its isogenic Δ*prmA* and Δ*relA* strains were grown in 5□mL TB7 (10□g/L tryptone buffered at pH 7.0 with 100□mM potassium phosphate) at 37 °C aerated at 225 rpm for 16 h. Cells were harvested by centrifugation at 9,400□×□g, washed once with PBS and stored at -80 °C for proteomics analysis. For additional experiments, *E. coli* BW22511 or BL21(DE3) and their isogenic derivatives (Δ*prmA*, Δ*relAΔspoT*, and Δ*prmA*Δ*relA*) were grown overnight in LB (10 g/L tryptone, 5 g/L yeast extract, 5 g/L NaCl) at 37 °C aerated at 225 rpm.

For (p)ppGpp experiments, the strains were washed once with M9 minimal medium (1x M9 salts, 3 g/L ferric citrate, 0.1 M CaCl_2_, 1 M MgSO_4_) and subcultured to an OD_600_ of 0.05 in 25 mL M9 salts supplemented with 0.2% casamino acids and 0.4% glucose.

### Quantitative proteomic analysis

*E. coli* proteins from the wild-type and Δ*prmA* strains were extracted, digested with trypsin, and analyzed by LC-MS/MS on Q-Exactive mass spectrometry (Thermo Fisher Scientific), as described in detail (26). LC-MS/MS data was processed with MaxQuant v.2.2.0.0, identifying peptides by search tandem mass spectra against the *E. coli* K12 sequences from Uniprot Knowledgebase downloaded on December 19, 2022. The search parameters included fully tryptic digestion with 2 missed cleavage sites, oxidation of methionine and lysine tri-methylation as variable modifications, and precursor mass tolerance of 20 and 4.5 ppm for first and main searches, respectively. Label-free quantification was set as default parameters and the “matching between runs” function was enabled using a matching window of 3 min and an alignment window of 20 min. Significantly changed proteins were determined by Student’s *t*-test.

### Polysome profiling

Polysome profiling was performed as described by Feid *et al*. (27). Briefly, 50 μL chloramphenicol (100□mg/mL) were added to every 50 mL of *E. coli* culture and cells were harvested and lysed in 10□mM Tris-HCl (pH 8.0), 10□mM MgCl_2_, and 1□mg/mL lysozyme by three freeze-thaw cycles. After lysis, 15 μL 10% sodium deoxycholate was added to the lysate, which was cleared by centrifugation at 9,400□×□g for 10 min at 4°C. A gradient of 10%-to-40% sucrose was prepared with BioComp Gradient Master model 108 in a buffer consisting of 20□mM Tris-HCl (pH 7.8), 10□mM MgCl_2_, 100□mM NH_4_Cl, and 2□mM dithiothreitol. Gradients were spun using an SW-41 rotor in an ultracentrifuge at 175,117□×□g for 3 h 45□min at 4°C. Fractions were collected using the ISCO/Brandel fractionation system by injecting a 50% sucrose solution below the gradient at 1.5□mL/min and monitored by absorbance at 254□nm.

### Measurement of cellular (p)ppGpp levels

Levels of (p)ppGpp were measured using a riboswitch Broccoli aptamer developed by Sun et al. (28); this aptamer binds to the probe DFHBI-1T and emits strong fluorescence in the presence of (p)ppGpp. Starved and control cells were incubated with DFHBI-1T and imaged using a Leica AF7000 Wide-field Fluorescence Microscope, using an APO PH3 100x/1.40 objective. To induce the biosensor, overnight cultures were washed once with M9 minimal medium then subcultured in M9 supplemented with 0.2% casamino acids and 0.4% glucose to an OD_600_ of 0.05. Cultures were grown at 37 °C aerated shaking at 225 rpm until an OD_600_ of 0.4 was reached. The biosensor and T7 polymerase were induced with 1 mM IPTG for 2 h. After 2 h, control samples were collected for DFHBI-1T incubation. The remaining culture was spun down and washed once with M9 salts then resuspended in 25 mL M9 salts. Cultures were starved for 15 minutes at 37 °C with aeration by shaking at 225 rpm. To measure fluorescence, 1 mL of the control or starved cultures was incubated with 200 μM DFHBI-1T at RT for 30 minutes. 10 μL of sample was spotted onto a glass slide and coved with a poly-L-lysine coated coverslip for imaging.

## Results

### Methylated proteomes of bacteria

To obtain a global landscape of the methylated lysine and arginine proteomes of bacteria, we searched a large phyloproteomics data comprised of 48 different bacterial strains from 6 different phyla for mono-, di- and tri-methylation sites (21). On average, 389, 136 and 123 proteins per bacterium were mono-, di- and trimethylated, respectively, (**Figure 1A**). In *Rhodopseudomonas palustris*, 904, 206 and 194 mono-, di- and tri-methylated proteins were identified (**Figure 1A**), showing the depth of methylated proteome coverage. Overall, methylated proteins represented a large portion of the total; on average, 23%, 8% and 7% of the identified cellular proteins were respectively modified by mono-, di- and tri-methyl groups (**Table S1**). In some extreme cases, such as *Prevotella ruminicola*, 39% of the identified cellular proteins were methylated (**Table S1**). These data show that methylation modifies a significant portion of the bacterial proteome across different species.

**Figure 1.**
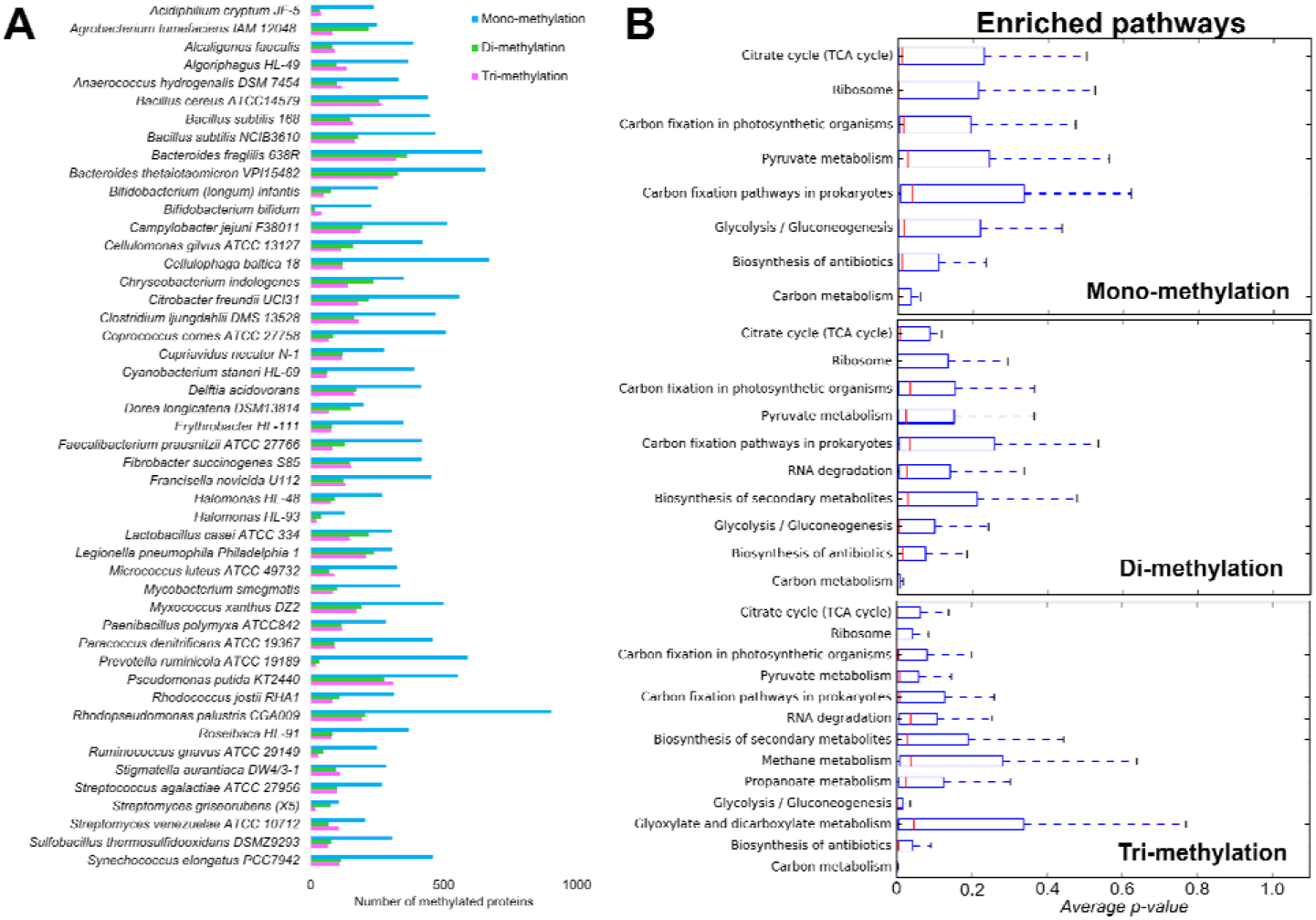
Phyloproteomics analysis of protein lysine and arginine methylation in bacteria. (A) Mono-, di-, and tri-methylated sites in 48 bacterial strains across the phylogenetic spectrum. (B) The most enriched pathways in these 48 strains.

We next investigated the cellular pathways enriched in methylated proteins across different bacterial strains. Several pathways were enriched in mono-, di-, and tri-methylated proteins, including central carbon metabolism (TCA cycle, carbon fixation, glycolysis/gluconeogenesis, and carbon metabolism), metabolism of antibiotics and secondary metabolites, ribosomes, and RNA degradation (**Figure 1B**). In addition, tri-methylated proteins were also enriched in proteins from glyoxylate, dicarboxylate, methane, and propane metabolism (**Figure 1B**). These data show that methylation modifies proteins from a variety of conserved pathways in bacteria.

We analyzed conserved methylation sites on individual proteins. The only methylation sites conserved throughout the phyloproteomics dataset were on proteins associated with translational machinery. This analysis showed that methylation is more conserved on the protein subunits close to the translation site of the ribosome (shown by the presence of mRNA and tRNA); bL11 and EF-Tu were among the subunits with the most conserved methylation sites (**Figure 2**). Sequence alignment of bL11 revealed conservation of these methylated sites (**Figure 3**). Overall, the analysis of the phyloproteomics dataset demonstrated that lysine and arginine methylation is a common post-translational modification in bacteria, with conserved sites in the translational machinery.

**Figure 2.**
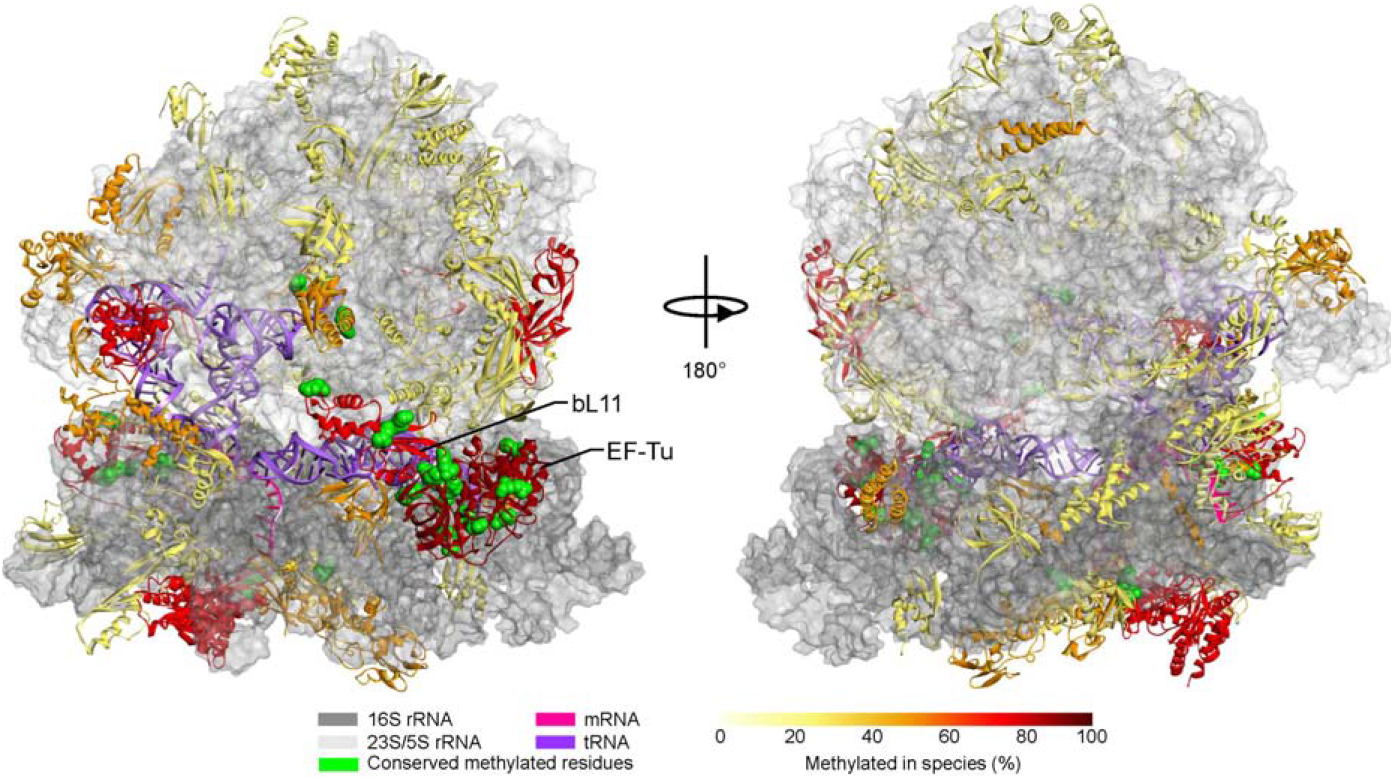
Conserved methylation sites in the translational machinery. Ribosomal proteins and elongation factor Tu 2 (EF-Tu) are represented by ribbon structure. Conserved methylated residues are in green in Corey–Pauling–Koltun representation. Surface representation of ribosomal RNA, 5S, 16S, and 23S, is gray.

**Figure 3.**
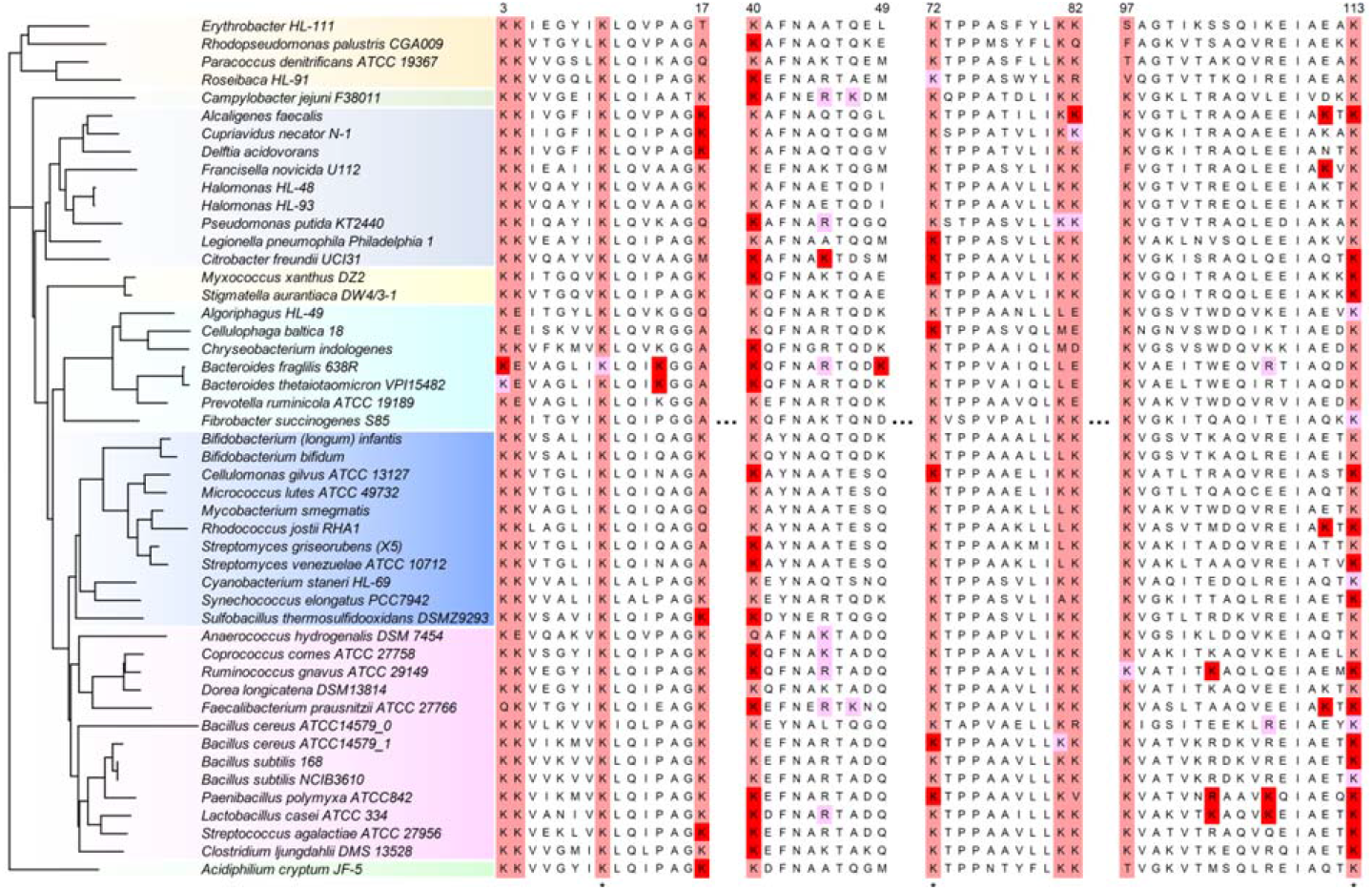
Methylated sites on bL11 in multiple sequence alignment. Species with conserved lysine greater than 60% are highlighted by light red. Tri-methylated residues and mono-/di-methylated residues are highlighted by red and pink, respectively. The asterisk at the bottom indicates 100% conserved lysines. Residue numbers are based on the bL11 sequence from *Escherichia coli* K-12.

### Proteomic analysis reveals abnormal regulation of cellular metabolism in a ΔprmA mutant during stationary phase

To determine whether protein methylation of ribosomal proteins plays a role in translation, we deleted the gene that encodes ribosomal protein L11 methyltransferase (*prmA*) in *E. coli* BW25113 and performed a proteomics analysis. As a previous report showed that a Δ*prmA* mutant of *Thermus thermophilus* exhibits no growth defect (10), we decided to investigate in the stationary phase and thus incubated our cultures for 16 h. Tri-methylation of lysines 3 and 4 of bL11 were only detected in the wild-type strain: these were the only methylations altered in the Δ*prmA* mutant. Furthermore, the unmodified version of the methylated peptide was only detected in the Δ*prmA* mutant, showing the modification stoichiometry to be 100% (**Figure 4A**). Our proteomic analysis identified and quantified 1772 proteins, with the expression of 272 proteins downregulated and 265 proteins upregulated in the Δ*prmA* mutant as compared to the wild type (**Figure 4B, Table S1**). On these proteins, we performed a functional-enrichment analysis using DAVID (Database for Annotation, Visualization, and Integrated Discovery) (29). Processes usually downregulated during cell starvation by the stringent response, such as translation (ribosome) and nucleotide metabolism (30), were upregulated in the Δ*prmA* mutant (**Figure 4C**). Pathways that are usually upregulated during cell starvation by the stringent response, such as alanine, aspartate and glutamate metabolism, pyruvate metabolism, and glyoxylate and dicarboxylate metabolism (30), were downregulated in the Δ*prmA* mutant (**Figure 4C**). These results indicate an abnormal regulation of cellular metabolism during stationary phase for the Δ*prmA* mutant.

**Figure 4.**
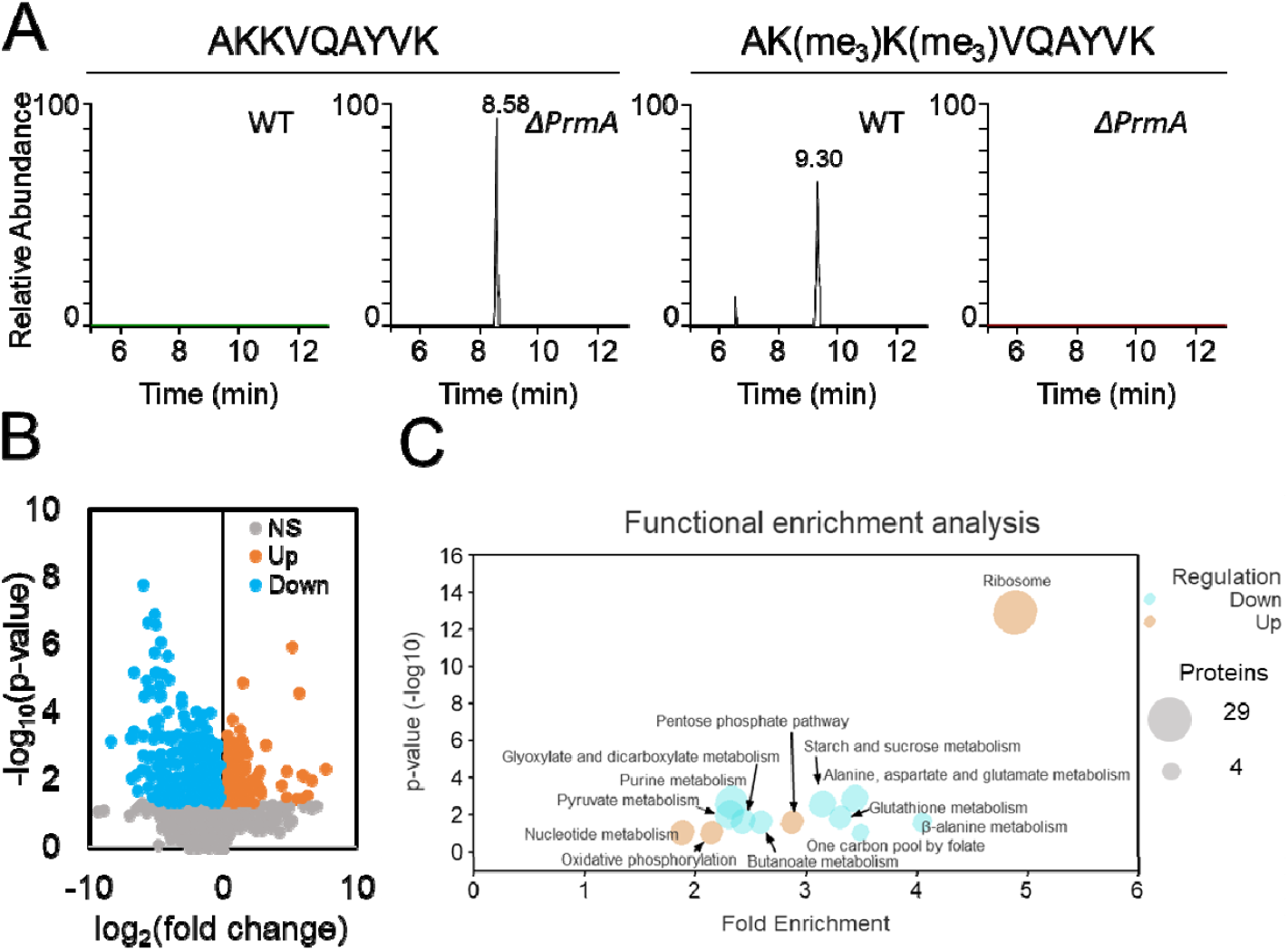
Proteomics analysis of the bL11 methyltransferase PrmA mutant strain. (A) Extracted-ion chromatogram of the bL11 N-terminus peptide containing tri-methylation on lysines 3 and 4. The modification is detected in the wild-type strain (n=3), but not in the Δ*prmA* mutant (n=4). (B) Differentially abundant proteins in the Δ*prmA* mutant versus the wild-type strain. (C) Functional-enrichment analysis using DAVID of the differentially abundant proteins identified in the proteomics analysis. The bubble plot shows the fold enrichment versus the statistical significance of each pathway. The size of the bubbles corresponds to the number of proteins matched to that specific pathway, while the colors indicated whether the pathway was enriched with up-(orange) or down-regulated (blue) proteins.

### bL11 trimethylation regulates the stringent response

During stationary phase, the stringent response is a major regulator of metabolism. Thus, we examined the proteomics data for several proteins involved in this response. We found that both enzymes producing the alarmone (p)ppGpp, RelA and SpoT, as well as the (p)ppGpp degradation enzyme GppA, were downregulated in the Δ*prmA* mutant (**Figure 5A**). (p)ppGpp usually reduces the levels of RNA in cells, but RNA polymerase subunits RpoA (beta) and RpoZ (omega) were upregulated in the Δ*prmA* mutant, as was DksA, which works with (p)ppGpp to downregulate transcription from strong promoters, especially those that transcribe genes involved in translation (**Figure 5A**). Conversely, the RNA degradation proteins RapZ and Rng were downregulated in the Δ*prmA* mutant (**Figure 5A**). Both effects indicated a possible increase in RNA concentration in cells, which we confirmed by extracting and quantifying RNAs (**Figure 5B**). (p)ppGpp also downregulates DNA replication, and the proteomics analysis revealed an increase in the DNA replication protein PriB (**Figure 5A**). (p)ppGpp also induces universal stress proteins (Usp) via RpoS, an alternative sigma factor that is activated during stationary phase. Levels of RpoS are not regulated by (p)ppGpp and, as such, was present at similar abundances in the mutant and wild type strains (**Figure 5A**). However, UspA, C, D, E, F and G were all reduced in the Δ*prmA* mutant (**Figure 5A**). Overall, the proteomics analysis led us to hypothesize that the mutant was impaired in (p)ppGpp production. To test this hypothesis, we used a Broccoli riboswitch assay that fluoresces in the presence of (p)ppGpp. When cells were starved, (p)ppGpp was strongly induced in the wild type but not in the Δ*prmA* mutant (**Figure 5C**). As a positive control, we also analyzed the Δ*relA* mutant, which showed complete depletion of (p)ppGpp production (**Figure 5C**). These results confirm that bL11 trimethylation affects (p)ppGpp levels, and thus the stringent response.

**Figure 5.**
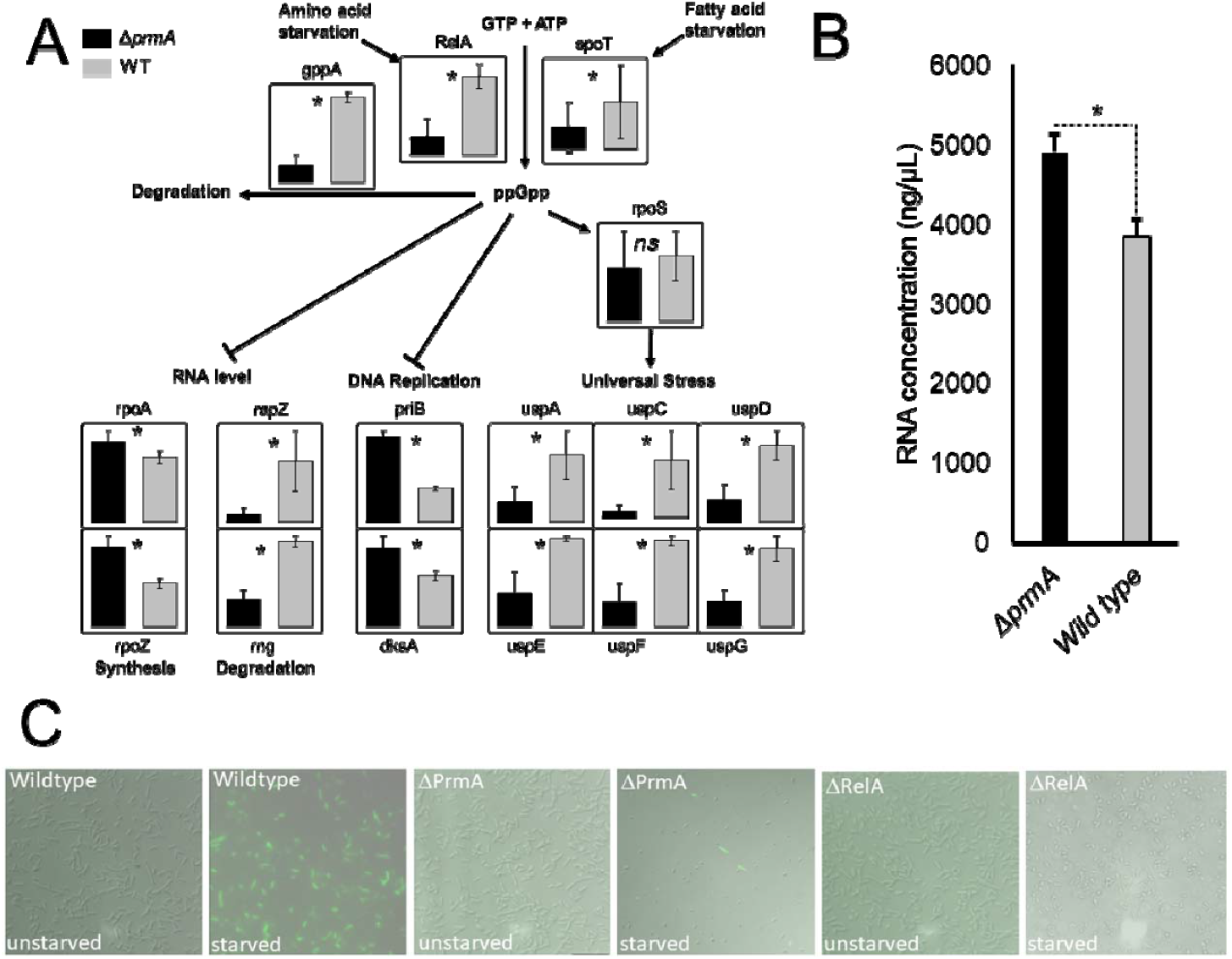
Regulation of the stringent response by bL11 methyltransferase PrmA. (A) Regulation of the stringent response proteins measured in the proteomics experiment. (B) RNA amounts in wild-type and Δ*prmA* strains. (C) Measurement of (p)ppGpp production by fluorescence microscopy. The green color is emitted by Broccoli riboswitch that fluoresces in the presence of (p)ppGpp. * p ≤ 0.05 by Student’s *t*-test.

### Lack of bL11 trimethylation leads to an abnormal polysome profile

In our proteomics analysis, we observed an increase in proteins associated with translation in the Δ*prmA* mutant relative to the wild type (**Figure 6A**). We thus examined the proteomics data for possible causes of this accumulation. Ribosomal proteins are degraded by the Lon protease, which is activated by the presence of polyphosphate (31). Although the levels of Lon did not significantly differ between the mutant and its parent, both polyphosphate kinase (PpK) and phosphatase (Ppx) were reduced in the Δ*prmA* mutant (**Figure 6B**), indicating lower production of polyphosphate. This lower level of polyphosphate would reduce Lon protease activity and thus promote the accumulation of ribosomes. Therefore, we performed a polysome profiling experiment to determine if the Δ*prmA* mutant would cause additional abnormality in the translational machinery. While making cell lysates, a white precipitate was observed in the Δ*prmA* mutant samples but not in the wild-type samples (**Figure 6C**). The precipitate was resistant to DNAse, RNAse and proteinase K (**Figure 6C**), indicating that DNA, RNA, or proteins might not be the major components of this precipitate or that they are somehow protected. We next performed polysome profiling of the Δ*prmA* mutant, comparing it to that of a Δ*relA* mutant. The Δ*relA* mutant in an unstarved condition exhibited a normal profile with most of the ribosomes in either the 70S form or as polysomes (**Figure 6D**); both are associated with protein translation. Conversely, when the Δ*relA* mutant was starved, the 30S and 50S subunits were favored (**Figure 6D**). A similar skewing towards the 30S and 50S subunits was observed with the Δ*prmA* mutant, even without starvation (**Figure 6D**). These data show that bL11 tri-methylation plays a role in association of the subunits into active ribosomes, despite the lack of a growth defect by the mutant (**Figure 6E**).

**Figure 6.**
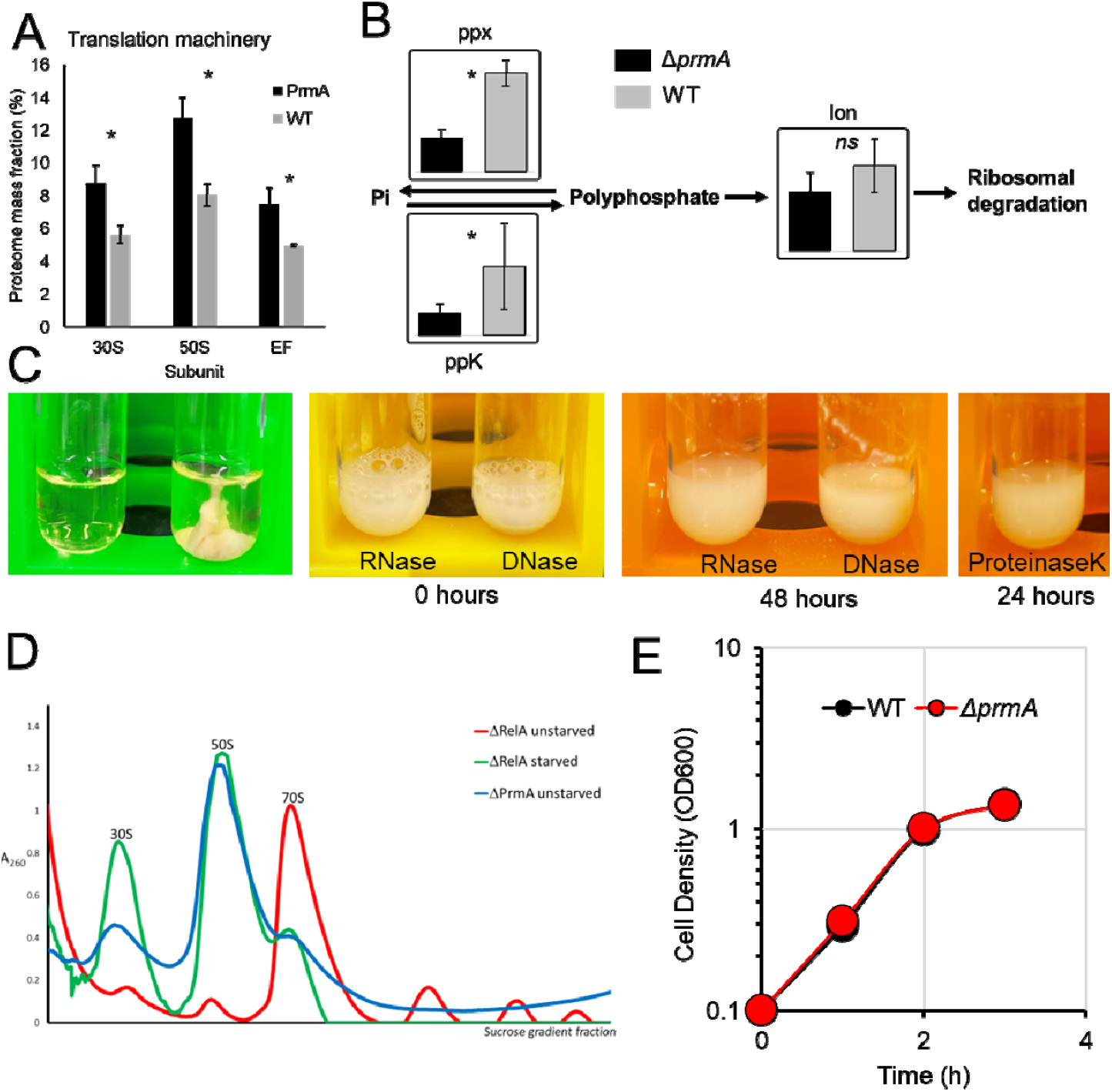
The effect of the Δ*prmA* mutant on ribosomes. (A) Levels of the translational machinery proteins measured in the proteomics analysis. (B) Levels of proteins from the polyphosphate pathway measured in the proteomics analysis. (C) Extracts of the wild-type and Δ*prmA* strains, and treatment of the latter with RNAse, DNAse and Proteinase K. (D) Polysomal profile of Δ*relA* and Δ*prmA* mutants in starved and unstarved conditions. (E) Growth curve of the wild-type and Δ*prmA* strains (n=6). * p ≤ 0.05 by Student’s *t*-test.

## Discussion

A recent review article by Macek and coworkers argued that lysine and arginine methylation are understudied PTMs in bacteria (2). Here, we show that approximately a quarter of the bacteria proteome is methylated with many conserved processes, including individual sites that are invariant across the bacterial evolution. Among the most conserved biological processes are central carbon metabolism and translation. In humans, the protein arginine methyltransferases PRMT5 and PRMT6 di-methylate enolase 1 at arginine 9, which enhances glycolytic activity and promotes cancer growth (32, 33). However, the roles of methylation in regulating the bacterial central carbon metabolism needs further investigation.

Regarding translation, we found that methylation is conserved on ribosomal protein subunits and the elongation factor Tu. Furthermore, we showed that trimethylations catalyzed by PrmA on bL11 conserved lysine residues are important for stringent response signaling and ribosomal homeostasis. The fact that bL11 trimethylation is important for the stringent response explains the lack of phenotype in exponentially growing *Thermus thermophilus* cells (10, 15) and reveals a function for this specific modification after almost half a century of study (14). Consistent with our findings, bL11 regulates RelA activity through its N-terminus, the region methylated (20). We also found bL11 methylation is important for ribosomal homeostasis in stationary phase. In this phase, part of the translational machinery is targeted for degradation to increase intracellular availability of nitrogen (34). Ribosomal degradation in stationary phase has been shown to be mediated by polyphosphate-enhanced Lon protease activity (31). However, how the stringent response increases the levels of polyphosphate has been a subject of discussion. Kornberg et al. showed that polyphosphate levels in stationary phase are increased by reducing the activity of polyphosphatase Ppx by (p)ppGpp (35), whereas Gray provided evidence that DksA enhances the expression of polyphosphate kinase Ppk (36). Our data show that bL11 methylation is also important for increasing Ppk levels, probably by regulating the levels of (p)ppGpp.

Methylation of ribosomal protein subunits promotes ribosomal biogenesis and maintains translation fidelity in the baker’s yeast *Saccharomyces cerevisiae* (37). Besides proteins, RNAs also get methylated, including messenger (mRNA), ribosomal (rRNA), non-coding (ncRNA) and transfer (tRNA) RNAs. The main types of RNA methylations are N6-adenosine (m^6^A) and 5-cytosine (m^5^C) methylations, which largely promote stabilization of transcripts in bacteria, archaea, and eukaryotes (38-40). Methylation of rRNAs help cells withstand stress conditions (39), while methylation of tRNAs promotes codon-anticodon interaction and prevents amino acid misincorporation (41). Methylation of mRNAs promotes the translation of specific transcripts (40). Therefore, methylation of several components of the translational machinery has functions in protein production.

In summary, our phyloproteomics analysis showed that a significant portion of bacterial proteins are methylated, including evolutionary conserved sites. bL11 trimethylation was among the most conserved methylation sites and we showed that this methylation is important for both stringent response signaling and maintenance of ribosomal homeostasis.

## Data Availability

Results of methylated peptide identifications in the phyloproteomics data were uploaded into the Open Science Framework at https://osf.io/4zpjh/. The proteomics data of the *E. coli* Δ*prmA* mutant versus wild-type strain are available at MassIVE under accession number MSV000092270.

## Acknowledgements

The authors thank Drs. Jared Shaw (University of Nebraska) and Nick Noinaj (Purdue University) for insightful discussions. Work was performed in the Environmental Molecular Science Laboratory, a U.S. Department of Energy (DOE) national scientific user facility at Pacific Northwest National Laboratory (PNNL) in Richland, WA. Battelle operates PNNL for the DOE under contract DE-AC05-76RLO01830.

## Author contributions

ESN, SHP, AJF and CVR proposed and designed the project; HEW, YF, MCB, SCF and DC performed the experiments; All the authors analyzed the data; ESN, SHP, AJW and CVR contributed with reagents and resources; ESN, HEW, SCF, AJW, YY, and CVR wrote the manuscript. All the authors edited and approved the final version of the manuscript.

